# Normative modelling of cortical networks identifies subtypes of type 2 diabetes associated with distinct clinical and transcriptomic profiles

**DOI:** 10.64898/2026.08.23.746498

**Authors:** Xiaoyue Wang, Zhizhong Sun, Jiancang Cao, Xinyuan Liang, Lianglong Sun, Jing Tian, Mingrui Xia, Lianping Zhao, Shijun Qiu, Yong He

## Abstract

Type 2 diabetes (T2D) is characterized by substantial clinical heterogeneity, yet its neurobiological substrates remain unclear. Here, we leverage normative modelling of cortical morphometric networks to quantify individual-level brain deviations using structural MRI data from 3,723 participants (1,941 T2D patients and 1,782 healthy controls) across three independent cohorts. Data-driven clustering reveals two robust T2D biotypes characterized by widespread positive (biotype 1) and negative (biotype 2) brain deviations. Despite comparable overall metabolic burdens, these biotypes feature distinct brain–metabolic coupling patterns: biotype 1 is primarily associated with lipid metabolism, whereas biotype 2 reflects a multifactorial metabolic burden spanning glycaemic control, adiposity, lipid, vascular risks, and disease duration. These divergent brain vulnerability profiles have distinct cognitive consequences; notably, biotype 2 shows poorer performance in complex cognitive tasks, aligning with negatively deviated connectivity gradients along the sensorimotor-to-association axis. Spatial transcriptomic analyses further link biotype 2 deviations to gene expression patterns enriched in insulin signalling, mitochondrial functions, tight junctions, and neurodegenerative disease pathways, alongside specific involvement of inhibitory neurons and oligodendrocyte lineage cells. Our findings provide a neurobiological understanding of T2D heterogeneity and establish a data-driven framework for characterizing personalized brain vulnerability, with implications for advancing precision diabetes medicine.

## Introduction

Type 2 diabetes (T2D) affects more than 500 million individuals globally ^1^ and is a major risk factor for accelerated cognitive decline and dementia ^2,3^. Although the clinical heterogeneity of T2D is well documented—with patients showing distinct disease trajectories, symptom severity levels, and complication risks ^4–7^—the neurobiological basis of this variability remains largely undefined. Structural neuroimaging studies have revealed T2D-associated brain alterations ^8–12^; however, the findings are notably inconsistent, ranging from widespread cortical thinning or volumetric reductions ^11,13,14^ to no detectable differences relative to healthy controls (HCs) ^15,16^. These discrepancies likely reflect substantial interindividual variability in the neuroanatomical signatures of the disease. Resolving this issue requires moving beyond treating T2D as a uniform entity and towards identifying neurobiologically coherent subtypes (or biotypes) that can explain the observed clinical heterogeneity and guide personalized interventions.

Previous efforts to delineate the neuroanatomical architecture of T2D have faced two key methodological challenges. First, the traditional case control designs inherently obscure the individual-level variance that is essential for precision phenotyping ^8–11^. While the recently developed normative brain charting addresses this issue by quantifying individual structural deviations against age- and sex-adjusted reference trajectories ^17–20^, this approach has not yet been applied to parse T2D heterogeneity. Second, most neuroimaging studies involving T2D focus on isolated, univariate morphological metrics ^14,16,21,22^, overlooking the highly coupled, network-level structural covariance that reflects axonal connectivity, cytoarchitecture, and gene expressions ^23,24^. Individualized morphometric similarity networks—such as those built using the morphometric inverse divergence (MIND) method ^25–27^—capture these multivariate features at the single-subject level. Integrating morphometric networks with normative modelling has proven effective for evaluating the phenotypic heterogeneity exhibited by individuals with neuropsychiatric conditions ^28–30^; however, this strategy remains unexplored for metabolic diseases such as T2D.

Here, we aggregated large-scale structural magnetic resonance imaging (MRI) data derived from 1,941 patients with T2D and 1,782 HCs. Utilizing a normative model of cortical morphometric networks derived from an independent lifespan cohort consisting of 33,937 participants (aged 0–80 years) ^30^, we quantified individualized network deviation scores against the typical age- and sex-adjusted trajectories. We clustered these individual-level deviations to stratify patients into distinct neuroanatomical biotypes and further evaluated their relevance to clinical and cognitive profiles. Finally, because specific genetic factors drive the phenotypic variance of T2D ^31–33^, we mapped these biotype-specific macroscopic brain deviation patterns onto spatial transcriptomic profiles to explore their molecular correlates. By defining ageing-calibrated, network-based, and molecularly informed T2D biotypes, we aim to provide a data-driven precision neurology foundation for metabolic diseases.

## Results

Following standardized processing and rigorous quality control schemes ^30^, we included high-quality structural MRI data derived from 1,941 patients with T2D (aged 27–84 years) and 1,782 HCs (aged 29–84 years) across three cohorts: the First Affiliated Hospital of Guangzhou University of Chinese Medicine (GZUCM) cohort (463 T2D patients and 260 HCs; 1 site), the Gansu Provincial Hospital (GPH) cohort (140 T2D patients and 107 HCs; 1 site), and the UK Biobank (UKB) cohort (1,338 T2D patients and 1,415 HCs; 4 sites) (Table S1). For each participant, the cortical surface was parcellated into 318 regions via a modified Desikan–Killiany atlas ^34^ (referred to as DK-318). Five morphological features—the surface area, cortical thickness, grey matter volume, mean curvature, and sulcal depth—were extracted for each vertex within these regions (Fig. 1a). These features were subsequently utilized to construct individual-level morphometric similarity networks via the MIND method (as detailed in the Methods section) ^27^. We computed regional morphometric similarity strength (MSS), which was defined as the average morphometric similarity of a given region to all others, and applied a normative model ^30^ to quantify a participant-specific deviation score for each region. We identified T2D biotypes by performing K-means clustering on the patients’ deviation maps (Fig. 1a) and evaluated their associations with clinical and cognitive characteristics (Fig. 1b), as well as transcriptomic signatures (Fig. 1c).

**Fig. 1.**
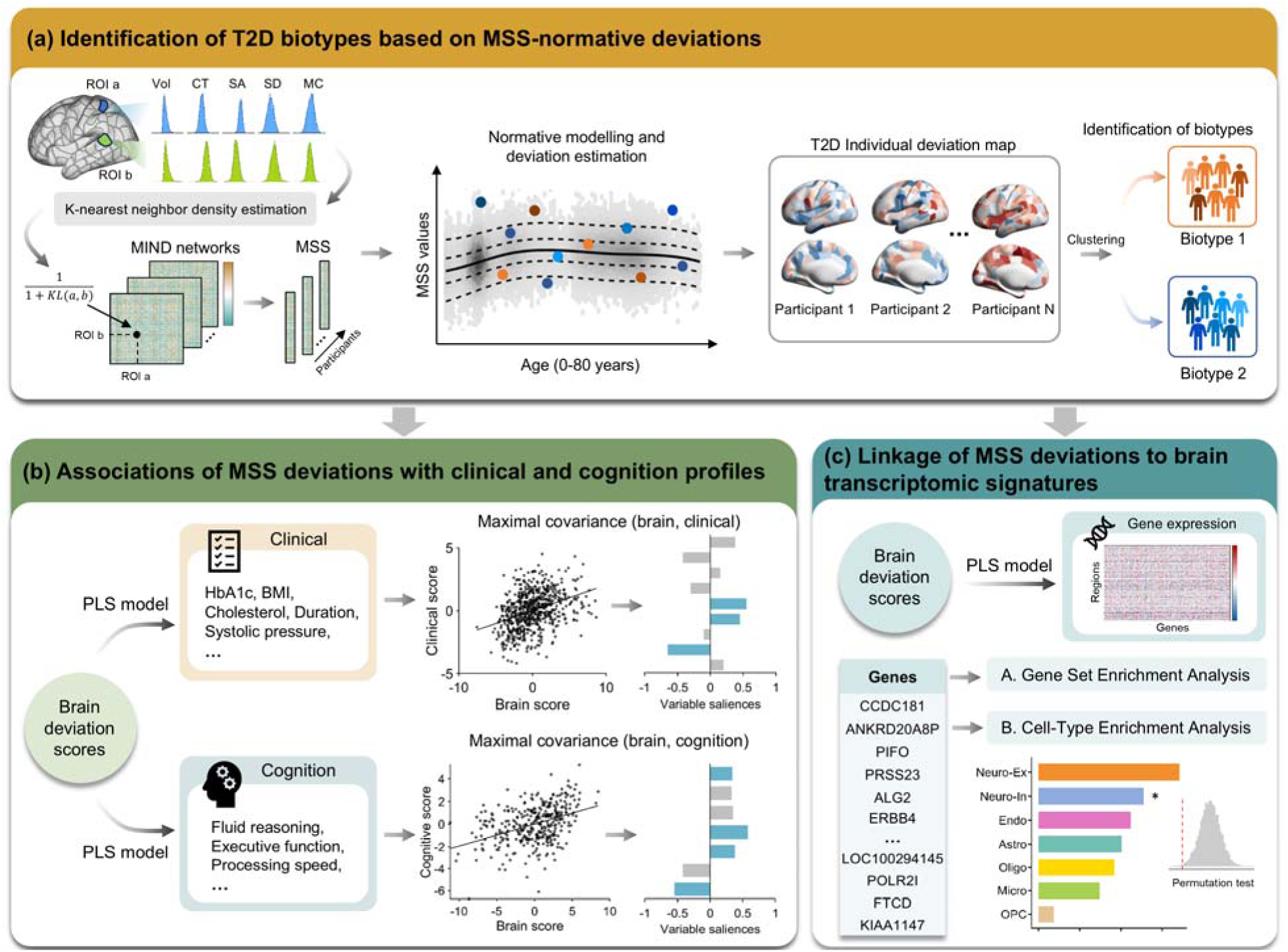
Flowchart of the study design. **(a)** Utilizing the MIND method, we constructed individualized morphometric similarity networks based on five morphological features ^27^ and derived regional MSS value for each participant. The established normative models ^30^ were subsequently used to quantify the regional MSS deviation scores, on the basis of which the T2D patients were stratified into distinct biotypes. **(b)** Within each identified biotype, a partial least-squares (PLS) correlation analysis was employed to characterize the multivariate coupling patterns between the brain deviations and multidimensional clinical profiles (encompassing clinical metabolic indicators and cognitive functions); this was followed by the identification of the core features driving these spatial associations. **(c)** The PLS regression framework was applied to spatially couple the biotype-specific deviation maps with the transcriptomic profiles using postmortem gene expression data from the Allen Human Brain Atlas ^35^, followed by a downstream gene set enrichment analysis ^36^ and a cell-type enrichment analysis. Cortical maps were visualized using BrainNet Viewer ^37^. SA, surface area; CT, cortical thickness; Vol, grey matter volume; MC, mean curvature; SD, sulcal depth; MSS, morphometric similarity strength.

### Morphometric network charts revealed substantial individual variability in T2D patients

To quantify the individual-level deviations of cortical morphological networks, we employed a normative modelling framework based on a generalized additive model for location, scale, and shape (GAMLSS) ^38^, built on structural MRI data of 33,937 participants (aged 0–80 years). In this framework, the MSS value of a specific brain region was set as the dependent variable, age was employed as a smoothing term, sex and the Euler number were utilized as covariates, and the scanning site was used as a random effect. For the three independent datasets (GZUCM, GPH, and UKB) that were not included in the original normative model ^30^, we implemented a rigorous out-of-sample estimation strategy ^39^. Specifically, to eliminate batch effects across the different MRI scanning sites, we used maximum likelihood estimation to align the HCs data of each site to the corresponding age span of the normative trajectories, estimating the site-specific statistical offsets (i.e., random effect intercepts). After updating the global normative model with these site-specific offsets, we applied the adjusted model to the entire cohort (including both T2D patients and HCs) of the corresponding site. The model then calculated the site-adjusted centile scores for each subject across all regions, which were subsequently converted into standard normal Z scores using the inverse cumulative distribution function. This process was iteratively performed across all 318 regions, ultimately generating a whole-brain deviation score matrix that precisely quantified the extent to which each individual’s MSS deviated from the norm.

Compared with the HCs, the patients with T2D had a greater total number of brain regions with extreme MSS deviations (|z| > 2), including both overall (*p* < 0.001) and positive (*p* < 0.01) deviations (Fig. 2a). Notably, although most patients had at least one extreme positive (76.9%) and negative (88.5%) deviation (Fig. 2b), no single region was highly deviated in more than 5.1% of patients (Fig. 2c). These results remained robust under a stricter threshold of |z| > 2.6 (Fig. S1). Together, these findings revealed substantial interindividual variability in the T2D-related cortical network alterations.

**Fig. 2.**
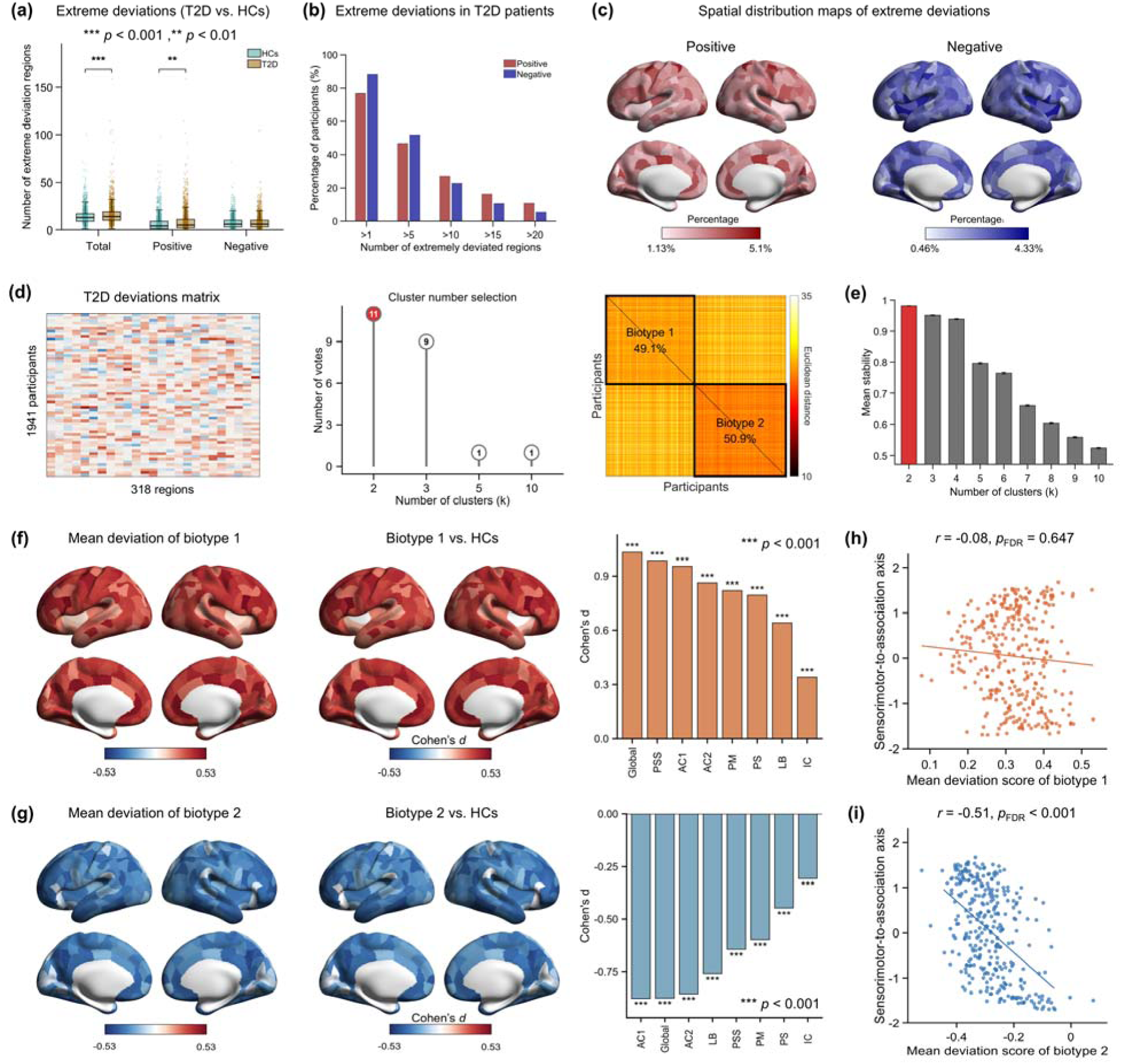
Normative modelling of cortical networks revealed the interindividual heterogeneity and biotypes of T2D patients. **(a)** The between-group differences in the number of extreme deviations (|z| > 2) between the patients with T2D and HCs. **(b)** The distribution of the number of extreme positive and negative deviated regions per T2D patient. **(c)** Spatial distribution maps showing the percentage of T2D patients with extreme deviations in each region. **(d)** Determination of the optimal number of T2D biotypes using the NbClust package (middle) ^43^. The heatmap visualizes the pairwise similarities among the MSS deviation patterns of T2D individuals, revealing two biotype clusters. **(e)** The robustness of the clustering structure was evaluated using a resampling-based clustering stability analysis. **(f, g)** Spatial distributions of the MSS deviation pattern of each biotype and their multilevel comparisons with the HCs. **(h, i)** Spatial associations between the mean MSS deviation pattern of each biotype and the sensorimotor-to-association axis ^41^. Statistical significance was evaluated via a spin test ^42^ and corrected for multiple comparisons via the FDR procedure. PS, primary sensory; PSS, primary/secondary sensory; PM, primary motor; AC1/AC2, association cortices; IC, insular; LB, limbic.

### Identifying robust T2D biotypes using individual deviation profiles of cortical network

By performing k-means clustering on the individual MSS deviation profiles for the patients with T2D (N = 1,941), we identified two T2D biotypes with distinct network-level alterations (Fig. 2d; detailed demographics are shown in Fig. S2). A resampling-based clustering stability analysis confirmed the robustness of this solution (mean stability = 0.98, 1,000 iterations; Fig. 2e).

Specifically, biotype 1 (N = 953, ages: 27–84 years) exhibited widespread positive MSS deviations, primarily involving the somatomotor, visual, prefrontal, and anterior and posterior cingulate cortices (Fig. 2f). Compared with the HCs, these patients had higher deviation scores globally, across all seven cytoarchitectonic classes ^40^, and in nearly all regions (317 regions with *p*_FDR_ < 0.05, false discovery rate-corrected; Cohen’s *d*: 0.10 to 0.53) (Fig. 2f). In contrast, biotype 2 (N = 988, ages: 28–84 years) showed widespread negative MSS deviations, primarily in the anterior and posterior cingulate, as well as the lateral frontal, parietal, and temporal cortices (Fig. 2g). The deviation scores for this biotype were lower than those of the HCs globally, across all seven cytoarchitectonic classes ^40^, and in over 96% of regions (307 regions with *p*_FDR_ < 0.05; Cohen’s *d*: -0.50 to -0.09) (Fig. 2g).

To further evaluate the spatial organization structure of these biotype-specific alterations, we examined the topographical correspondence between the mean deviation pattern of each biotype and the cortical sensorimotor-to-association axis ^41^. This analysis revealed a significant alignment for biotype 2, which was assessed via a permutation test that accounted for spatial autocorrelation (spin test) ^42^ (*r* = -0.51, *p*_spin_ < 0.0001, *p*_FDR_ < 0.001; 10,000 iterations; Fig. 2i). Specifically, the magnitudes of the negative MSS deviations became progressively more pronounced from the primary sensorimotor regions towards the high-order association cortices. In contrast, the deviation pattern of biotype 1 was not significantly associated with this cortical axis (*r* = -0.08, *p*_spin_ = 0.647, *p*_FDR_ = 0.647; Fig. 2h).

### Symptom correlates of T2D biotype-specific cortical network deviations

Before examining the clinical profiles of the identified biotypes, we first analysed their baseline demographic and lifestyle characteristics. The results revealed that the patients in the biotype 2 group were significantly older than those in the biotype 1 group (*p* < 0.001, Cohen’s *d* = -0.30) and were more likely to be male (*p* = 0.001, odds ratio = 1.35) (Figs. S2b–c). In contrast, no significant group differences were detected across the key lifestyle factors, including smoking, alcohol consumption, and physical activity (Supplementary Results; Table S2), suggesting that the observed biotype divergence is unlikely to be primarily explained by these behavioural habits. Accordingly, age, sex, and site were included as covariates in all subsequent analyses to account for these baseline demographic differences.

After controlling for these baseline demographic factors, we next examined the biotype-specific differences among the clinical profiles using a univariate analysis of covariance (ANCOVA) (Fig. S3, Table S3). Compared with the HCs, both T2D biotypes showed clinical and metabolic phenotypic alterations, including elevated body mass index (BMI), glycated haemoglobin (HbA1c), fasting plasma glucose (FPG), and triglyceride (TG) (all *p*_FDR_ values < 0.01). Compared with biotype 2, biotype 1 had higher systolic blood pressure (SBP) (*p*_FDR_ = 0.020). The two biotypes did not significantly differ in BMI, disease duration, glycaemic control (HbA1c and FPG), homoeostasis model assessment 2 of insulin resistance (HOMA2IR), homoeostasis model assessment 2 of β-cell function (HOMA2B), or lipid profiles (total cholesterol (TC), TG, high-density lipoprotein (HDL), and low-density lipoprotein (LDL)).

We employed multivariate partial least-squares (PLS) analyses to investigate whether these comparable clinical–metabolic profiles were coupled with brain deviations in a biotype-specific manner. The multivariate analyses revealed distinct brain–metabolic coupling patterns between the two biotypes. For biotype 1, the multivariate PLS analysis revealed that the first latent variable (LV1) accounted for 28.1% of the shared covariance between the brain deviation and clinical metabolic profiles (*p*_perm_ = 0.03, permutation test 10,000 iterations; Fig. 3a), with a significant correlation observed between the latent brain scores and the clinical scores (*r* = 0.32, *p* < 0.001; Fig. 3b). To further delineate the specific features driving this covariation pattern, the stable contributions of the variables were evaluated. The results indicated that the clinical indicators (HDL, TC, and LDL) that were closely tied to lipid metabolism exhibited stable positive loadings (Fig. 3c). Correspondingly, the brain regions with stable contributions manifested almost exclusively as widespread negative loadings, which were predominantly concentrated in the inferior parietal, posterior cingulate, temporo-parieto-occipital junction, sensorimotor, and inferior frontal cortices (Fig. 3d). Mapping these significant regions onto the seven cytoarchitectonic classes ^40^ further revealed that they are concentrated mainly in the primary motor and multimodal association cortices (Fig. 3d).

**Fig. 3.**
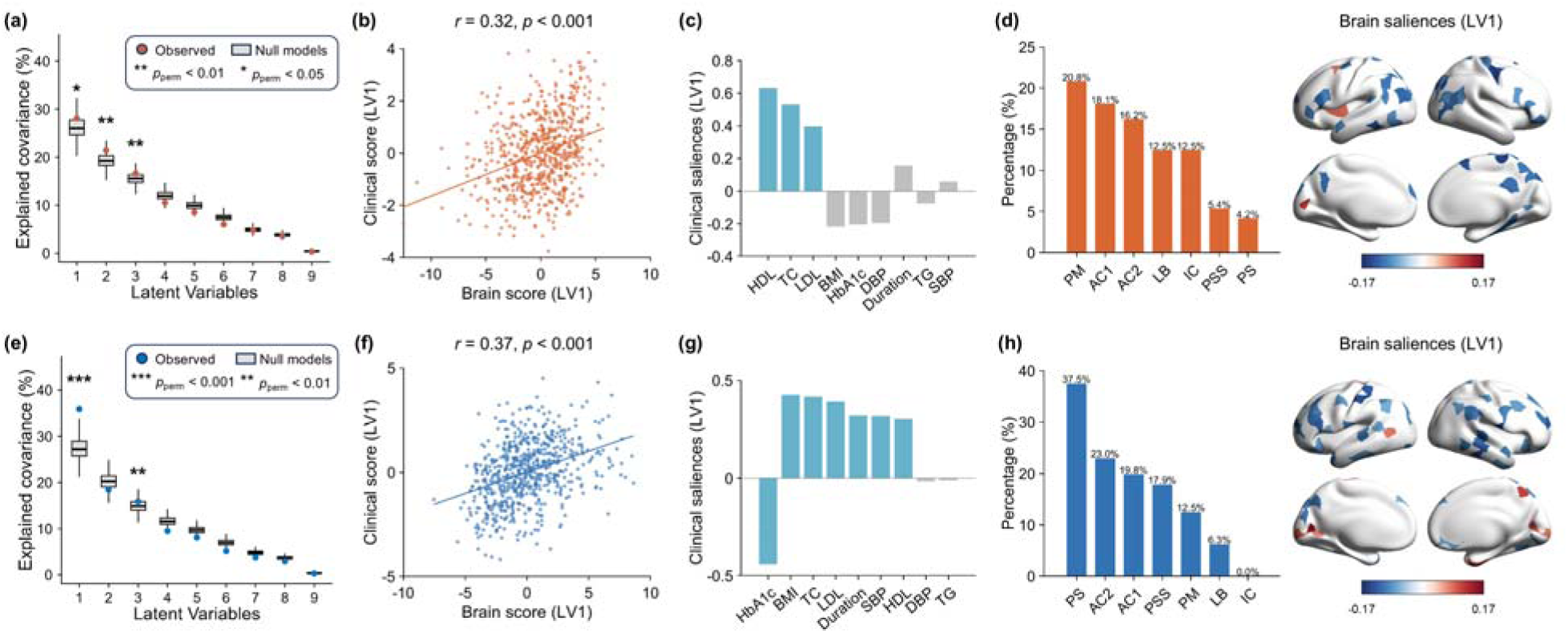
Biotype-specific multivariate brain–clinical associations. (a–d) Results derived for biotype 1 and **(e–h)** results derived for biotype 2 from PLS correlation analyses. **(a, e)** Percentage of the explained covariance between the brain deviation scores and clinical scores captured by each LV. Boxplots represent the null distributions generated from 10,000 permutations, with dots denoting the empirical values. **(b, f)** Pearson correlation between the brain score and clinical score within LV1. **(c, g)** Contribution weight of each clinical metric to LV1, where the stability of these weights was assessed using bootstrap resampling (1,000 iterations). Colour-coded bars indicate variables that exceeded the bootstrap ratio (BSR) threshold (|BSR| > 2). **(d, h)** Bar plots (left) showing the percentage of stably contributing regions within each cytoarchitectonic class ^40^. Cortical maps show the regions with stable contributions (|BSR| > 2) to LV1 (right). HDL, high-density lipoprotein; TC, total cholesterol; LDL, low-density lipoprotein; BMI, body mass index; HbA1c, glycated haemoglobin; DBP, diastolic blood pressure; TG, triglyceride; SBP, systolic blood pressure.

For biotype 2, the multivariate PLS analysis revealed that LV1 accounted for 35.9% of the shared covariance between the brain deviation and clinical metabolic profiles (*p*_perm_ < 0.001, 10,000 iterations; Fig. 3e), with a significant correlation observed between the latent brain scores and the clinical scores (*r* = 0.37, *p* < 0.001; Fig. 3f). An evaluation of the clinical variables driving the PLS coupling pattern revealed that HbA1c, BMI, TC, LDL, the disease duration, SBP, and HDL exhibited stable contributions (Fig. 3g). These features collectively indicated a broader systemic metabolic axis encompassing glycaemic dysregulation, adiposity, lipid metabolism, the vascular burden, and the disease duration. The brain regions with stable contributions were similarly dominated by widespread negative loadings and were predominantly distributed across the sensorimotor, dorsolateral prefrontal, lateral temporal and visual cortices (Fig. 3h). Mapping these stably contributing regions onto the seven cytoarchitectonic classes ^40^ revealed that they are concentrated mainly in the primary sensory cortex, followed by the multimodal association cortices and primary/secondary sensory areas (Fig. 3h).

### Cognitive correlates of T2D biotype-specific cortical network deviations

To investigate whether the two T2D biotypes exhibited distinct cognitive phenotypes, we conducted a behavioural performance evaluation across four major cognitive domains: processing speed (trail-making #1 and symbol digit substitution), executive function (trail-making #2), memory (including working memory assessed through numeric memory and episodic memory assessed by pairs matching), and fluid reasoning (fluid intelligence and matrix pattern completion) (detailed in Table S4).

Utilizing an ANCOVA adjusted for age, sex, education level, and site, we found significant group-level differences across all the assessed measures (all *p*_FDR_ values < 0.05). These results showed that compared with the HCs, both biotypes exhibited poorer cognitive performance (Fig. 4a), yet they differed substantially in both the specific domains affected and the deficit degrees. Post hoc pairwise comparisons revealed that biotype 1 displayed mild-to-moderate, relatively restricted cognitive impairments, scoring significantly lower than the HCs did, primarily in specific tasks such as trail-making #1, symbol digit substitution, numeric memory, and matrix pattern completion. In contrast, biotype 2 exhibited pervasive and severe cognitive impairment. Notably, the patients in biotype 2 performed significantly worse than both the HCs and the biotype 1 patients did in tasks including trail-making #2, fluid intelligence, and matrix pattern completion, highlighting a profound cognitive vulnerability that was specific to this biotype. To determine whether the pronounced cognitive deficits observed for biotype 2 were merely a consequence of a longer disease duration, we performed a nonparametric Mann Whitney U test, which revealed no significant diabetes duration difference between the two biotypes (*p* = 0.492; Fig. S4b). Furthermore, the observed cognitive deficits in the biotype 2 remained robust despite these patients being significantly older than those in the biotype 1 group were (*p* < 0.001; Fig. S4a).

**Fig. 4.**
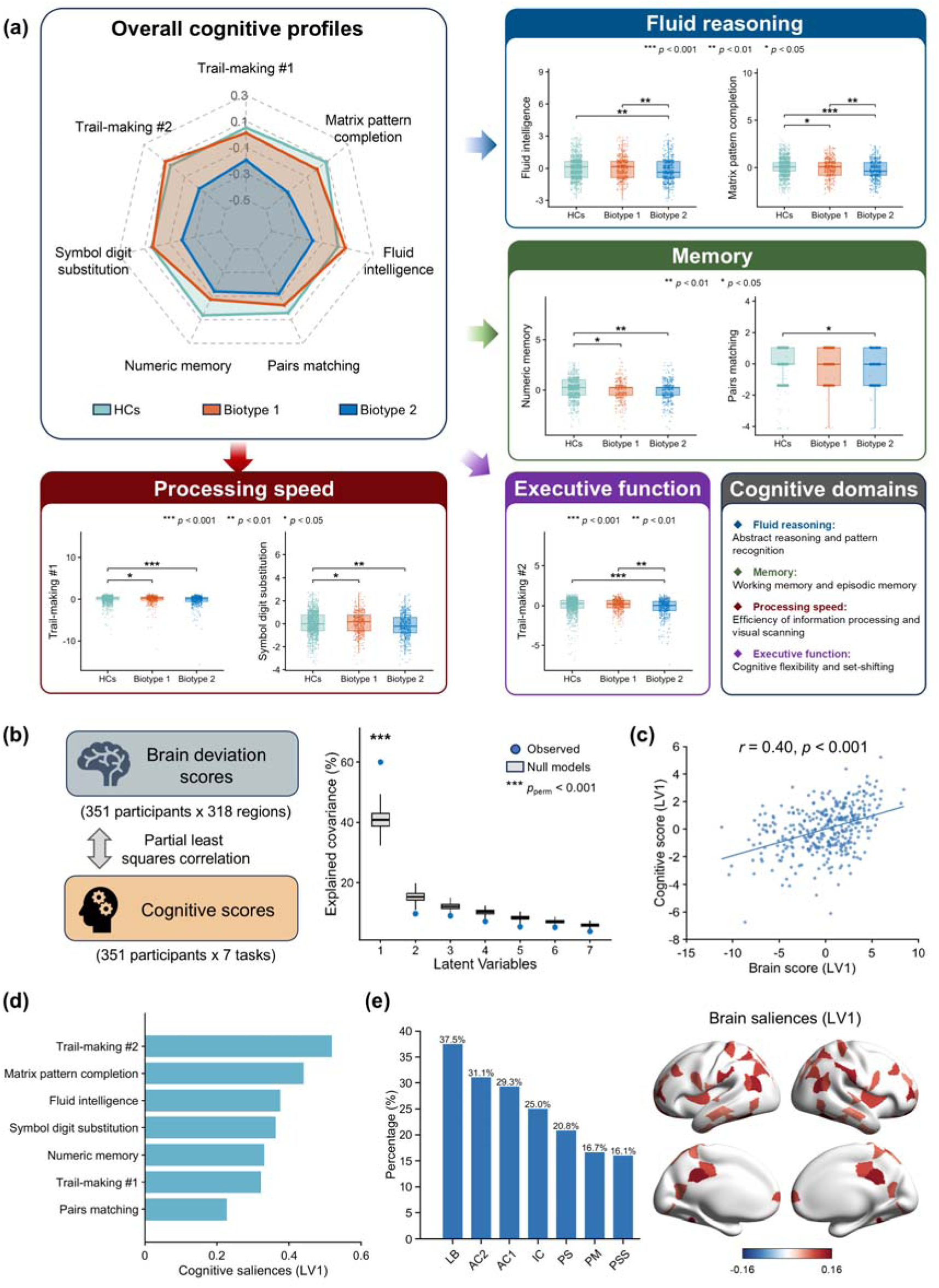
Cognitive profiles and multivariate brain-cognition associations produced across both biotypes. Cognitive performance was assessed in a subset of the UKB cohort (1352 HCs, 580 patients with biotype 1, and 627 patients with biotype 2). **(a)** Radar plots summarizing the cognitive performance across the seven tasks for the HCs and biotype 1 and biotype 2 individuals. Boxplots show the group differences observed for each cognitive task. **(b–e)** PLS correlation analysis of the brain-cognition associations of biotype 2. **(b)** Percentage of explained covariance between the brain deviation scores and the cognitive scores captured by each LV (right). Boxplots represent the null distributions generated from 10,000 permutations, with blue dots denoting the empirical values. **(c)** Pearson correlation between the brain score and the cognitive score for the latent space of LV1. **(d)** Contribution weight of each cognitive metric to LV1, where weight stability was assessed using bootstrap resampling (1,000 iterations); colour-coded bars indicate variables exceeding the stability threshold (|BSR| > 2). **(e)** The bar plot (left) shows the percentage of these robust regions relative to the total number of regions contained within each cytoarchitectonic class ^40^. The cortical map shows the spatial distribution of the regions with stable contributions (|BSR| > 2) to LV1 (right).

A multivariate PLS analysis revealed that only biotype 2 showed significant associations between brain deviations and cognitive profiles. LV1 explained 60% of the cross-covariance between the brain and cognitive measures in the biotype 2 patients (*p*_perm_ < 0.001, 10,000 iterations; Fig. 4b), and there was a significant correlation between the resulting brain scores and cognitive scores (*r* = 0.40, *p*_perm_ < 0.001; Fig. 4c). Within the LV1 dimension, all the cognitive metrics and brain regions yielded widespread and stable positive contributions (Fig. 4d). These brain regions were concentrated mainly in the limbic and multimodal association cortices and involved multiple areas, such as the dorsolateral prefrontal, medial prefrontal, parietal, posterior cingulate, and temporal cortices (Fig. 4e). This pattern indicates that a greater negative whole-brain deviation magnitude is correlated with more severe cognitive impairments across multiple domains.

### Transcriptomic signatures of T2D biotype-specific cortical network deviations

To elucidate the molecular mechanisms driving these biotype-specific morphological network deviations, we spatially mapped the brain deviation patterns of each biotype onto transcriptomic data derived from the Allen Human Brain Atlas (AHBA) ^35^. The results of a PLS regression analysis revealed that only the deviation pattern of biotype 2 exhibited a significant spatial–transcriptomic association. Specifically, the first PLS component (PLS1) explained 20.38% of the variance in the biotype 2 deviation pattern (*p*_spin_ = 0.033, spin test: 10,000 iterations; Fig. 5b), and the regional PLS1 gene score was significantly correlated with the brain deviation pattern (*r* = 0.45, *p*_spin_ = 0.033; Fig. 5c). Spatially, these regional scores were relatively elevated in the lateral temporal, posterior cingulate, and medial prefrontal cortices and the paracentral lobule, whereas lower scores were observed in the somatosensory, visual, insular, and temporo-parieto-occipital junction cortices (Fig. 5c).

**Fig. 5.**
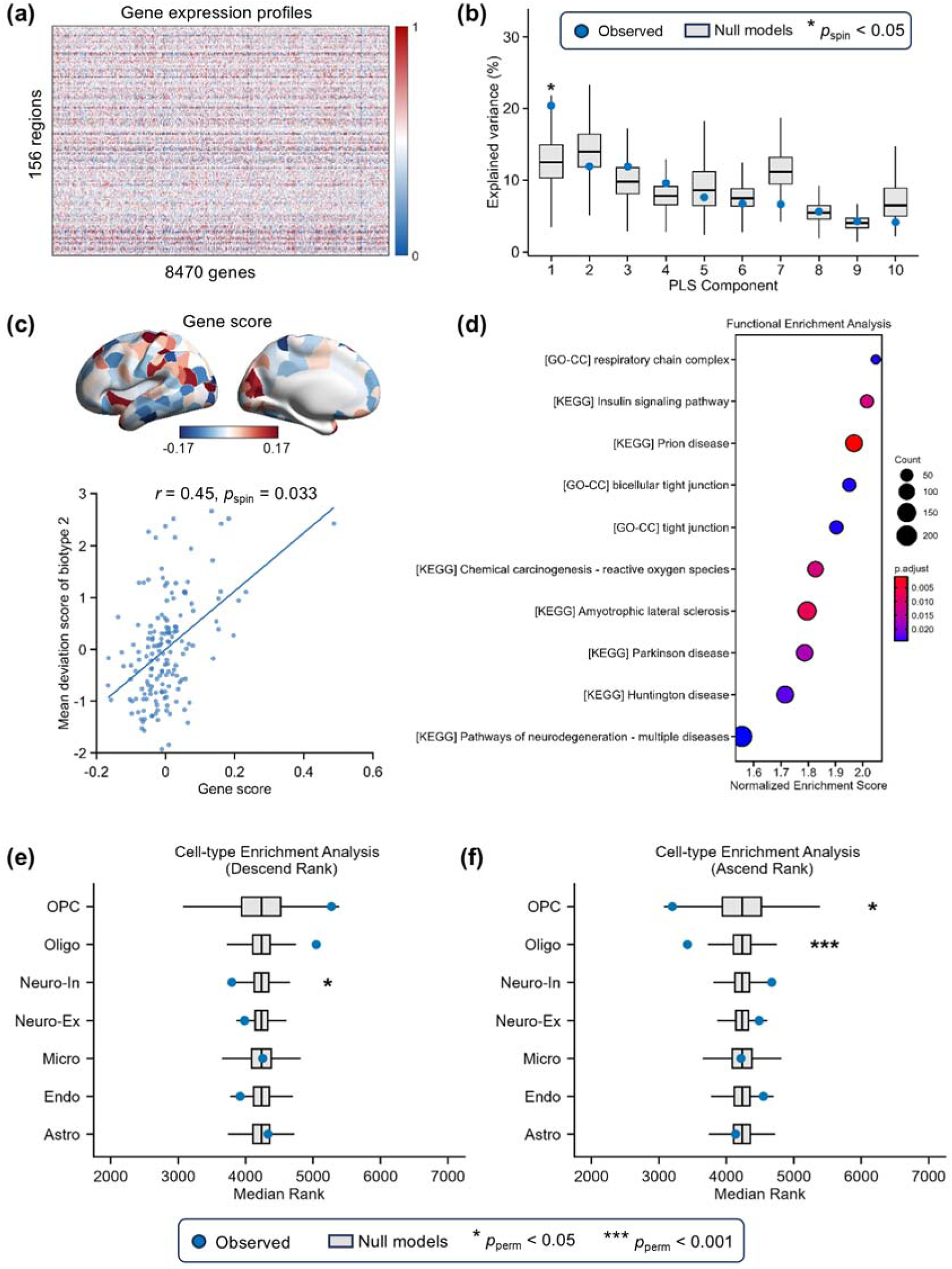
Biotype 2-specific transcriptomic correlates and cell-type enrichment of brain deviations. **(a)** Gene expression profiles across different brain regions. **(b)** Percentage of variance explained by each PLS component. Boxplots represent the null distributions generated from the spin test (10,000 iterations), with the dots indicating the empirical values. **(c)** Distribution of the PLS1 gene scores and the spatial correlation between the gene scores and mean deviation scores for biotype 2, with statistical significance evaluated via a spin test. **(d)** Bubble plot illustrating the significantly enriched KEGG pathways and GO-CC terms based on the gene list ranked according to PLS1 weights in descending order. **(e, f)** Cell-type enrichment results obtained based on the gene lists sorted by the PLS1 weights in **(e)** descending and **(f)** ascending order. For each of the seven distinct cell types ^44^, the empirical median rank of its marker genes within the ranked list is denoted by a dot. The boxplots represent the null distributions of the median ranks generated from 10,000 random permutations. KEGG, Kyoto Encyclopedia of Genes and Genomes; GO-CC, Gene Ontology Cellular Components; Micro, microglia; Endo, endothelial cells; OPC, oligodendrocyte precursor cells; Oligo, oligodendrocytes; Astro, astrocytes; Neuron-ex, excitatory neurons; Neuron-in, inhibitory neurons.

To further characterize the biological underpinnings of this transcriptomic pattern, we calculated individual gene weights for the PLS1 component and evaluated their stability using a bootstrapping procedure. We then performed a gene set enrichment analysis (GSEA) ^36^ on the full gene list, which was sorted in descending order according to the bootstrap-derived z scores. A Kyoto Encyclopedia of Genes and Genomes (KEGG)-based pathway enrichment analysis revealed that genes at the top of this ranked list (reflecting the strongest positive weights) were significantly enriched in the insulin signalling pathway, chemical carcinogenesis (reactive oxygen species), various neurodegenerative disease pathways (including prion disease, amyotrophic lateral sclerosis, Parkinson’s disease, and Huntington’s disease) and pathways of neurodegeneration (multiple diseases) (Fig. 5d, Table S5). Concurrently, a gene ontology (GO) enrichment analysis further revealed three significant cellular component terms, namely, the respiratory chain complex, bicellular tight junction and tight junction, which are related mainly to mitochondrial energy metabolism and intercellular tight junctions (Fig. 5d, Table S5). At the cellular level, a cell-type enrichment analysis conducted based on the seven canonical cell classes defined by Seidlitz et al. ^44^ revealed cell-type-specific associations with the biotype 2 spatial deviation pattern. Specifically, the marker genes for inhibitory neurons (Neuron-in) were significantly enriched among the genes with high positive weights (*p*_FDR_ = 0.021; Fig. 5e), whereas the marker genes for oligodendrocytes (Oligo: *p*_FDR_ < 0.001) and oligodendrocyte precursors (OPC: *p*_FDR_ = 0.027) were significantly enriched the among genes with negative weights (Fig. 5f).

### Sensitivity analyses

A series of sensitivity analyses were conducted to confirm the stability of the identified T2D biotypes across multiple dimensions. First, a leave-one-site-out cross-validation demonstrated remarkably high spatial consistency between the rederived biotype patterns and the main findings across all iterations (biotype 1: all *r* values > 0.91; biotype 2: all *r* values > 0.92; all *p*_spin_ values < 0.001; Fig. S5a), indicating that the biotype architecture captures robust disease-related morphological patterns rather than site-specific biases. Second, despite the inherent demographic variations, repeated independent clustering processes performed on the UKB dataset (*r* = 0.89–0.95; all *p*_spin_ values < 0.001; Fig. S5d) and the combined Chinese cohorts (GZUCM and GPH) (*r* = 0.62–0.79; all *p*_spin_ values < 0.001; Fig. S5b) yielded spatial patterns that were significantly correlated with the main findings (Fig. S5c), highlighting the cross-population stability of these biotypes. Finally, to assess whether the biotyping results were sensitive to the strategy used for estimating site effects, we recalculated the deviation scores using an alternative normative modelling approach. Specifically, half of the newly introduced control data were incorporated into the training set to estimate site effects (Fig. S6a) ^38^. Under this approach, the optimal two-cluster solution was preserved, and the resulting deviation profiles were strongly correlated with the main findings (biotype 1: *r* = 0.93; biotype 2: *r* = 0.91; all *p*_spin_ values < 0.001;

Fig. S6b). Crucially, 96.75% of patients maintained consistent biotype assignments across the two deviation scoring frameworks (Fig. S6c). Furthermore, applying a modified Desikan–Killiany atlas comprising 219 regions ^30^ (referred to as DK-219) yielded identical biotype numbers and deviation profiles (Figs. S7a–b), with 93.97% individual-level assignment concordance with the DK-318 scheme (Fig. S7c). Importantly, a significant spatial correlation with the sensorimotor-association axis was found only for biotype 2 (*r* = -0.48, *p*_spin_ < 0.001, *p*_FDR_ < 0.001; Fig. S7d). Collectively, these analyses support the robustness of the T2D neurophysiological biotypes across different imaging sites, population subsets, normative calibration strategies, cortical parcellation schemas, and lifestyle-related factors.

## Discussion

In the present study, we utilized normative modelling to elucidate the neurobiological heterogeneity among patients with T2D, identifying two distinct neuroimaging-derived biotypes that were characterized by widespread positive (biotype 1) and negative (biotype 2) morphometric deviations. Although the two biotypes exhibited similar clinical metabolic profiles, they showed distinct brain–metabolic coupling patterns, with biotype 1 primarily associated with lipid metabolism and biotype 2 reflecting a broader multifactorial metabolic burden. Notably, biotype 2 exhibited a greater vulnerability profile, which was characterized by poorer cognitive performance and distinct molecular and cellular signatures that were associated with its spatial deviation pattern. Together, these findings reveal a neurobiological dimension of T2D heterogeneity beyond the conventional metabolic characterization and highlight the value of neuroimaging-based biotyping for understanding diabetes-related brain vulnerabilities.

Although biotype 1 and biotype 2 demonstrated clear differences between their macroscopic brain structures, their baseline peripheral metabolic profiles, including HbA1c, FPG, insulin resistance indices, and disease durations, were largely comparable. These findings suggest that diabetes-related brain alterations are unlikely to be explained solely by the severity of peripheral metabolic disturbances. Rather, under similar metabolic burdens, interindividual differences in the intrinsic response patterns of the brain or in the inherent resilience to metabolic insults may play a more critical role. Although no significant differences were observed in the individual clinical–metabolic metrics between the two biotypes, a multivariate PLS correlation analysis revealed distinct, biotype-specific coupling patterns between the metabolic profiles and brain deviations. In biotype 1, brain deviations were predominantly associated with lipid metabolic indices (TC, LDL, and HDL). Such a lipid-centric coupling pattern may reflect neurovascular dysfunction and low-grade neuroinflammatory processes that are driven predominantly by dyslipidaemia. In contrast, biotype 2 exhibited a widespread and multifactorial-driven coupling pattern, in which glucose metabolism (HbA1c) constituted the dominant contributor, which was accompanied by significant contributions from the BMI, the disease duration, lipid profiles, and blood pressure. This multidimensional coupling suggests that negative brain deviations are jointly shaped by multiple interacting metabolic disturbances rather than by a single metabolic pathway. Previous studies have indicated that multiple comorbidities accompanying T2D, such as hyperglycaemia, dyslipidaemia, and hypertension, may have complex destructive effects on brain functions through diverse mechanisms ^2,45,46^. The deep coupling of this systemic metabolic burden with the pervasive negative brain deviations observed for biotype 2 implies that the brain networks of this population exhibit extremely high vulnerability when coping with multiple metabolic stressors. Our findings generate hypotheses regarding potential biotype-specific intervention strategies: the predominant brain–lipid coupling in biotype 1 raises the possibility that lipid management and microvascular protection may be particularly relevant for neuroprotection in this subgroup, whereas the multifactorial coupling observed for biotype 2 suggests that a comprehensive approach that simultaneously addresses blood glucose, blood pressure, lipids, and body weight might be necessary. However, these implications remain speculative, as cross-sectional correlational evidence cannot be used to establish whether modifying these metabolic factors would alter brain structure trajectories. Prospective intervention trials stratified by neuroimaging-derived biotypes are warranted for testing these clinical hypotheses.

The widespread, multifactorial brain–metabolic coupling observed for biotype 2 suggests that the neural networks of this subgroup might face a more complex systemic burden, potentially rendering them more vulnerable to functional decline. Although both biotypes exhibited cognitive deficits, which was consistent with the findings of previous studies ^2,47–50^, they demonstrated distinct impairment patterns. Biotype 2 demonstrated broader and more pronounced impairments in complex tasks, such as fluid intelligence, executive functioning, and matrix pattern reasoning. This phenomenon may be linked to its neuroanatomical organization scheme: the brain deviations of biotype 2 progressively intensified along the cortical sensorimotor-to-association axis, extending from the primary sensorimotor regions towards higher-order association cortices. Given the central role of association cortices in complex cognitive functions, this spatial gradient provides a plausible neuroanatomical context for the greater cognitive impairment observed for biotype 2. Consistent with this interpretation, significant multivariate brain-cognition coupling was observed only for biotype 2, with the stable brain contributions primarily involving widespread multimodal association cortices. Importantly, the more severe cognitive deficits observed for biotype 2 do not appear to be exclusively driven by a more advanced disease stage. The lack of a significant difference between the diabetes durations of the two biotypes does not support the hypothesis that they simply represent the early and late stages of T2D progression, suggesting instead that they may reflect fundamentally distinct neurobiological profiles within the T2D population. However, the patients with biotype 2 were significantly older, which might reflect a complex, synergistic destructive effect between T2D-based metabolic stress and normal brain ageing, ultimately compromising the resilience of higher-order neural networks. The classic work by Biessels et al. highlights that T2D-associated cognitive dysfunction is a highly heterogeneous continuum, spanning from subtle cognitive decrements to pervasive cognitive impairments ^2^. Our findings provide a neuroimaging-derived biotyping framework that maps onto this clinical continuum.

The relatively localized cognitive impairments in biotype 1 may represent a milder phenotype associated with basic metabolic dysfunction within this spectrum, whereas the extensive deviations of the higher-order association cortices in biotype 2 patients align it with the more severe end of the continuum. Although the cross-sectional nature of our study design prevented us from assessing whether biotype 2 patients will eventually develop clinically defined mild cognitive impairments or dementia, the pervasive structural alterations observed along the sensorimotor-to-association axis suggest a heightened vulnerability of this biotype when facing the dual pressures of metabolic stress and ageing, underscoring the clinical value of its early identification.

A spatial transcriptomic analysis further provided a molecular context for the brain deviation pattern of biotype 2. Genes whose spatial expressions covaried with regional damage were significantly enriched in the insulin signalling pathway, mitochondrial respiratory chain complexes, and intercellular tight junctions. Given that the AHBA reflects the baseline gene expression in neurotypical brains, these spatial alignments suggest inherent regional vulnerability rather than direct disease-induced alterations. Specifically, regions with inherently lower baseline expressions of these genes may possess fewer molecular reserves to buffer against systemic metabolic stress, rendering them more susceptible to structural disruptions. This hypothesis is biologically plausible and aligns with pathological evidence that chronic systemic metabolic disorders can significantly downregulate endothelial tight junction proteins and disrupt the integrity of the blood–brain barrier, which in turn induces central nervous system insulin resistance ^45,51–53^. Furthermore, the significant enrichment of reactive oxygen species-mediated oxidative stress and multiple classic neurodegenerative disease pathways—including prion disease, Parkinson’s disease, and amyotrophic lateral sclerosis—in our KEGG analysis reinforces the biological overlap between T2D and neurodegeneration. Overall, these findings lend imaging and transcriptomic support to the “Type 3 diabetes” and brain insulin resistance hypotheses, suggesting that T2D and neurodegenerative diseases might share underlying mechanisms of energy failure, mitochondrial dysfunction, and neurotoxicity ^45,53–55^. While causality cannot be inferred from spatial correlations, these molecular associations help explain the pervasive, neurodegenerative-like decline observed in biotype 2 patients. Future studies employing *in vivo* molecular imaging or transcriptomic analyses of patient-derived brain tissues are warranted to directly validate these pathway alterations and their causal contributions to the T2D-related neurodegeneration process.

A cell-type enrichment analysis further suggested that the biotype 2 deviation pattern was associated with specific cellular architectures. Specifically, the marker genes for oligodendrocytes and oligodendrocyte precursor cells were densely clustered at the extreme negative end of the gene list. This spatial correspondence suggests that anatomical hubs characterized by a high abundance of these myelinating cells might represent the primary regions associated with the structural vulnerabilities observed for biotype 2. Given the essential role of oligodendrocytes in myelin maintenance and their high energy demands, T2D-related insulin resistance and mitochondrial dysfunction may compromise their energy supply, potentially contributing to widespread myelin degradations and structural network collapses. This hypothesis aligns with cytological evidence showing that insulin signalling is crucial for the differentiation, maturation, and myelinating functions of oligodendrocyte lineages ^45,56,57^. In addition, the enrichment of inhibitory neurons suggests a potential role of excitation/inhibition (E/I) imbalances within local cortical microcircuits in aggravating metabolic stress and neurotoxicity; however, this speculation must be rigorously validated by future cell-level investigations.

Crucially, our brain-derived biotypes provide a critical and complementary neurophysiological dimension for classic clinical T2D subtyping ^5^. A cross-modal mapping analysis revealed that patients with similar clinical–metabolic phenotypes could still exhibit distinct brain injury patterns (Supplementary Results; Fig. S8), suggesting that relying solely on conventional metabolic indicators may not fully capture the neurophysiological heterogeneity exhibited by T2D, potentially delaying early neurological prevention and management processes.

Incorporating neuroimaging-derived normative models alongside traditional clinical classification schemes holds promise for enabling a comprehensive multimodal framework to characterize brain vulnerability, thereby facilitating precise neurological risk stratification and the future development of targeted neuroprotective therapies in patients with T2D.

Several limitations should be addressed. First, detailed treatment regimens (e.g., medication histories) and broader lifestyle or environmental exposures were not fully incorporated. Given that glucose-lowering drugs ^58,59^ and multidimensional lifestyle factors ^60–63^ profoundly influence both peripheral metabolism and neurocognitive health, their potential confounding effects on the observed biotype-specific brain deviation patterns and cognitive impairments could not be completely ruled out. Future studies controlling for these factors are essential for clarifying these impacts. Second, the present analysis focused exclusively on cortical morphometric similarity networks and excluded subcortical structures. Because the pathology of T2D also impacts deep subcortical nuclei (e.g., the hippocampus, amygdala, and thalamus) ^10,64–66^, future studies should incorporate cortico-subcortical systems to fully characterize T2D-related brain alterations and their cognitive relevance. Finally, the cross-sectional design limits the causal inferences regarding longitudinal disease trajectories. Future longitudinal cohorts are essential for evaluating the temporal stability of these biotypes. Given that biotype 2 may represent a high-risk prodromal phenotype for dementia, tracking its long-term clinical evolution trend is crucial for validating its prognostic value and facilitating early precision interventions.

## Methods

### Participants

This study included participants from three independent cohorts. These included two clinical cohorts in China, i.e., the GZUCM cohort and the GPH cohort, as well as a cohort sourced from the UKB. Each cohort consisted of patients with T2D and HCs. Following strict inclusion and exclusion criteria (detailed in the Supplementary Methods), the initial cohorts comprised 909 participants in the GZUCM cohort (597 T2D patients aged 27–74 years, with 373 males; 312 HCs aged 26–74 years, with 115 males), 266 in the GPH cohort (154 T2D patients aged 25–69 years, with 121 males; 112 HCs aged 29–66 years, with 62 males), and 3,356 in the UKB cohort (1,678 T2D patients aged 48–84 years, with 1014 males; 1,678 HCs aged 49–84 years, with 1013 males).

All study protocols and data collection procedures were conducted in strict accordance with the principles of the Declaration of Helsinki. Specifically, the GPH study received multistage ethical approval from the Medical Ethics Committee of Gansu Provincial Hospital (approval nos.: 2017-188, 2019-196, and 2023-500). The GZUCM study was approved by the Ethics Committee of the First Affiliated Hospital of Guangzhou University of Chinese Medicine (approval nos.: 2020-115 and 2023-146). The UKB study was approved by the North West Multi-centre Research Ethics Committee (approval nos.: 11/NW/0382, 16/NW/0274, and 21/NW/0157). Written informed consent was obtained from all participants prior to their enrolment and MRI acquisition across all three cohorts.

### Structural MRI data acquisition parameters

Structural MRI data for all participants were acquired using 3.0-Tesla (3.0T) MRI scanner. The specific acquisition parameters for each dataset are detailed as follows. (i) GZUCM cohort: Data were acquired using a 3.0 T Siemens MAGNETOM Prisma scanner equipped with a 64-channel head coil. The parameters were as follows: repetition time (TR) = 2530 ms; echo time (TE) = 2.98 ms; flip angle (FA) = 7°; field of view (FOV) = 256×224 mm^2^; matrix size = 256×224; and isotropic voxel size = 1.0×1.0×1.0 mm^3^. (ii) GPH cohort: Scanning was performed on a 3.0 T Siemens MAGNETOM Skyra scanner equipped with a 32-channel head coil. The parameters were as follows: TR = 2,530 ms; TE = 2.35 ms; FA = 7°; FOV = 256×256 mm^2^; matrix size = 256×256; and isotropic voxel size = 1.0×1.0×1.0 mm³. (iii) UKB cohort: Data were acquired on a 3.0 T Siemens MAGNETOM Skyra scanner equipped with a 32-channel head coil. The parameters were as follows: TR = 2000 ms; TE = 2.01 ms; FA = 8°; FOV = 256×256 mm^2^; matrix size = 256×256; and isotropic voxel size = 1.0×1.0×1.0 mm^3^.

### Structural data preprocessing

The structural data were preprocessed using the Human Connectome Project (HCP) structural pipeline (v4.4.0-rc-MOD-e7a6af9), which was containerized and deployed via the QuNex platform (v0.93.2). This preprocessing workflow encompassed three sequential stages. (a) The PreFreeSurfer stage involved applying brain extraction, resampling, spatial denoising, and bias field correction to the raw T1w structural data. (b) The FreeSurfer stage focused on the accurate reconstruction of cortical surfaces from the standardized structural images. Through the anatomical segmentation of brain tissues, high-resolution pial, white, and mid-thickness surfaces were generated, followed by spatial registration to the standard fsaverage surface atlas. (c) The PostFreeSurfer stage involved converting the processed data into the HCP standard format (CIFTI space). Volumetric data were aligned to the standard MNI space via nonlinear registration, and cortical topological surface data were mapped to the standard fs_LR_32k space through spherical registration and surface downsampling. Finally, to obtain regional morphological metrics at the individual level, the parcellation atlas in the standard fsaverage space was inversely registered back to each participant’s native anatomical surface space.

### Image quality control

To ensure the quality of the whole-brain geometric and topological reconstruction, a combination of automated assessment techniques and expert manual review was employed to comprehensively evaluate the quality of the structural MRI scans across all participants. The overall quality control (QC) workflow and exclusion criteria were strictly consistent with those of our previous study ^30^.

*(i) Automated raw image QC:* Prior to the preprocessing stage, the MRI quality control (MRIQC) tool was utilized to automatically extract image quality metrics (IQMs) from the raw T1w images. For each dataset, a participant was classified as an outlier and excluded if their structural image exceeded 1.5 times the global interquartile range (IQR) on at least three core IQMs. These six core automated QC metrics included the entropy focus criterion, the foreground-to-background energy ratio, the coefficient of joint variation, the contrast-to-noise ratio, the signal-to-noise ratio (SNR), and Dietrich’s SNR (SNRd).
*(ii) Topological surface quality assessment:* Upon the completion of surface reconstruction in the FreeSurfer stage, the Euler number was employed to quantitatively assess the geometric and topological complexity levels of the cortical surface, thereby identifying and excluding structural scans that were unsuitable for the subsequent analyses. A higher Euler number indicated fewer topological defects and superior cortical surface reconstruction quality. Participants were deemed to have surface reconstruction failures and were strictly excluded if their individual Euler numbers fell below 1.5 times the IQR of the study-specific distribution (Q1-1.5*IQR) or were less than -217.
*(iii) Manual visual inspection QC:* Following the automated QC screening procedure described above, five neuroimaging experts with anatomical expertise (X.Y.W., W.W.W., M.H.Z., C.Z., and M.L.L.) conducted a visual quality check on all the final structural scans. The visual QC criteria focused strictly on detecting artefacts and evaluating the quality of the cortical segmentation, surface reconstruction, and surface registration. Scans with quality defects were flagged by recording the participant identifiers, and X.Y.W. subsequently aggregated and double-checked all the visual inspection results to compile the final exclusion list.

Through these rigorous screening procedures, our final combined sample included 1,941 patients with T2D and 1,782 HCs with high-quality structural images for the subsequent analyses. Specifically, the final analyses included 463 T2D patients and 260 HCs from the GZUCM cohort, 140 T2D patients and 107 HCs from the GPH cohort, and 1,338 T2D patients and 1,415 HCs from the UKB cohort (Table S1).

### Construction of cortical morphometric networks

To construct individual cortical morphometric networks, the cerebral cortex was first parcellated into 318 cortical regions on the basis of a randomly modified Desikan–Killiany atlas ^34^, and individual morphometric networks were subsequently evaluated using the state-of-the-art MIND method ^27^. For each participant, five morphometric features—the cortical thickness, mean curvature, sulcal depth, surface area, and grey matter volume—were extracted for each vertex within each brain region to form a five-dimensional vector. Each feature was Z-standardized across all vertices to place the five features on a comparable scale. A Kullback–Leibler (KL) divergence estimator based on the k-nearest neighbours algorithm was then utilized to compute the multidimensional KL divergence between the feature distributions of any two brain regions.

Specifically, for each vertex i in brain region a (with a total of *v_a_* (*i*)vertices), we searched within the same region to find the k-th nearest neighbour of i and calculated their Euclidean distance in the five-dimensional feature space, with the result denoted as *p_k_* (*i*)Similarly, we searched all vertices in brain region b to find the k-th nearest neighbour of vertex i and calculated the Euclidean distance in the five-dimensional feature space, with the result denoted as *v_k_* (*i*)The unidirectional KL divergence from brain region a to b was defined as follows:

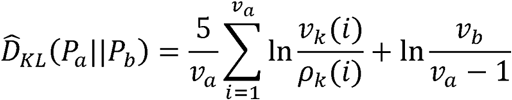

We also calculated the unidirectional KL divergence from region b to region a, which was denoted as *D̂_KL_*(*P_b_|P_a_*). The KL divergences from both directions were then combined for symmetrical reconstruction, yielding the final symmetrical KL divergence between brain regions a and b:

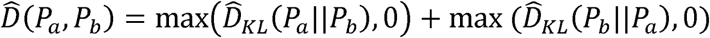

We applied a reciprocal transformation using the following formula to convert the divergence (representing dissimilarity) into a similarity measure. This transformation ensured that MIND = 1 when the feature distributions of two regions were perfectly identical, while MIND approached 0 when the divergence became infinitely large:

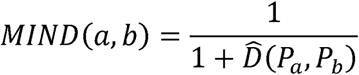

Following the aforementioned procedures, the individual MIND morphometric similarity network was constructed for each participant. Finally, the morphometric similarities between each brain region and all other regions were averaged to derive the MSS value for that region.

### Normative modelling and deviation score computation

In our previous study ^30^, a normative model was established for tracking the age-related trajectories of MSS values across each brain region. This model was trained on a large-scale dataset comprising more than 30,000 healthy individuals utilizing the GAMLSS framework ^38^ to derive the specific model parameters (*μ, σ, ν, τ*) for each brain region. We refer to this framework as the reference normative baseline, and none of the independent datasets utilized in the present study were included in the training cohort of this reference normative model.

Based on the parameterized structure of this normative model, we subsequently employed an out-of-sample estimation strategy ^39^ to map all new participants from the current cohorts onto the healthy baseline, thereby quantifying their individualized morphometric deviation scores.

Specifically, to eliminate systematic biases induced by multicentre scanners and imaging protocols, we first leveraged the HCs data within each specific site, along with the nonlinear developmental trajectories determined from the original reference norm, to estimate site-specific random effects. After correcting for these site effects, for any individual i in the new dataset, we extracted the individual’s age, sex, scanner site label, and Euler number. These individual metrics were then fitted into the updated GAMLSS normative architecture to derive the theoretical healthy population distribution values for the four morphometric probability parameters conditional on their specific covariate background: (*μ_i_, σ_i_, ν_i_, τ_i_*)

Subsequently, by integrating the empirical MSS values measured from each cortical region of that specific individual with the theoretically predicted parametric distribution derived from the normative baseline, their corresponding centile score was quantified via the cumulative distribution function. This centile score was further transformed into a standardized deviation score (Z score). The ultimately generated deviation score matrix represented the multiples of the standard deviation by which each individual’s MSS value deviated from the age- and sex-matched healthy normative baseline in each brain region.

### T2D biotyping based on brain-morphometric deviation features

To identify the interindividual heterogeneity of abnormal brain morphologies in patients with T2D, we performed an unsupervised k-means clustering analysis on the individualized deviation score matrix across 318 cortical regions for all patients. To select the optimal number of clusters within a candidate range from k = 2 to 10, the NbClust algorithm was utilized to evaluate the clustering quality across multiple dimensions ^43^. A total of 22 quantitative indices compatible with the k-means algorithm participated in the majority voting scheme, encompassing (i) geometric topology criteria balancing intracluster compactness and intercluster separation boundaries (e.g., the Calinski–Harabasz index, Davies–Bouldin index, Silhouette index, and Dunn index); (ii) matrix-derivative criteria tracking the first- or second-difference incremental inflections of the trace or determinant of sum-of-squares matrices across the multidimensional feature space as the k value increased (e.g., the Krzanowski–Lai index, Hartigan index, Scott index, Marriot index, and Tracew index); and (iii) statistical expectation criteria benchmarked against null random background distributions (e.g., the cubic clustering criterion index, McClain index, Ball index, and Ratkowsky index). The algorithm ultimately aggregated the independent votes from these 22 valid quantitative indices, employing a majority rule approach to determine the definitive optimal biotyping solution. To safeguard the stability and convergence of the partitioning results, the final k-means clustering process involved 10 independent random initializations and a maximum of 1,000 iterations in the background.

To further validate the robustness of the biotype structures derived from the unsupervised clustering, we performed a resampling-based clustering stability analysis. Specifically, a total of 1,000 independent resampling iterations were executed for each k value. During each iteration, two independent subsamples (denoted as subsample A and subsample B), each comprising 80% of the total T2D cohort, were generated via random sampling without replacement ^67^. The standard k-means clustering algorithm was then separately applied to subsamples A and B. To minimize the potential biases induced by the stochastic initialization of the cluster centroids, each individual clustering run was configured with 10 random initializations and a maximum of 1,000 iterations. Upon obtaining the two sets of clustering labels, we extracted the overlapping subjects common to both subsamples and recorded their respective biotype assignments. Owing to the inherent randomness of label numbering in unsupervised clustering, we addressed the label switching phenomenon by constructing a confusion matrix for the overlapping subjects and formulating the label alignment process as a linear assignment problem. The Hungarian algorithm was subsequently employed to identify the optimal bijective mapping that maximized the clustering consensus between the two partitioning solutions ^68^. Ultimately, the clustering stability for each individual iteration was quantified as the proportion of overlapping subjects who received identical partitioning assignments after conducting label alignment relative to the total number of overlapping subjects. After the 1,000 iterations were completed, the mean stability score was calculated for each k value, and the specific k yielding the highest mean stability was defined as the mathematically optimal and most robust biotyping solution.

To investigate whether the identified biotypes differed significantly in basic demographic characteristics, we performed statistical analyses for age and sex while consistently adjusting for the data collection sites to mitigate potential heterogeneity across the multicentre data. To evaluate age differences, a linear mixed-effects model was constructed with age as the dependent variable, biotype and sex as fixed effects, and the site as a random effect. Additionally, Cohen’s *d* was calculated to quantify the effect sizes between groups. For the analysis of the differences between the sex proportions, a logistic regression model was employed, where binarized sex was set as the dependent variable and biotype and site were included as independent variables. The significance levels for all between-group differences were derived from the main effect coefficients of the biotypes in their respective models.

### Characterization of biotype-specific morphological deviations

To verify whether the T2D biotypes identified via unsupervised clustering reflected genuine pathological characteristics that deviated from normative trajectories and to comprehensively profile their unique spatial deviation patterns, we conducted intergroup comparisons by benchmarking them against HCs across three topological scales: global, systemic, and regional. Intergroup statistical differences and effect sizes were quantitatively evaluated using the nonparametric Mann Whitney U test and Cohen’s *d*, respectively. At the global level, we compared the whole-brain averaged deviation scores between each biotype and HCs, further employing permutation tests with 10,000 iterations to guarantee the statistical robustness of the differences. At the systemic level, we parcellated the cerebral cortex into 7 cytoarchitectonic classes based on the von Economo atlas ^40^, compared the average deviation scores within each class, and applied the FDR method to correct for multiple comparisons. Finally, at the regional level, we strictly adhered to the aforementioned nonparametric testing and FDR correction pipeline to individually assess the intergroup differences across all cortical regions, thereby pinpointing the critical localized impairments driving the biotype-specific signatures.

### Statistical comparison of clinical and metabolic profiles across biotypes

To explore whether different neurophysiological biotypes exhibit varying degrees of disease progression and metabolic impairment, we evaluated the baseline clinical and metabolic differences across groups. Intergroup differences for the metabolic metrics shared among patients and HCs—namely, the BMI, HbA1c, SBP, diastolic blood pressure (DBP), TC, triglycerides (TG), HDL, LDL, FPG, and fasting insulin (FINS)—were assessed using an ANCOVA within the General Linear Model framework, following a sequential two-stage statistical procedure:

*(i) Determining the omnibus group effect via a nested model comparison:* To rigorously evaluate the overall clinical and metabolic discrepancies across groups while parsing out the variance of confounding factors, a nested model comparison approach was introduced. Specifically, a reduced model incorporating only the nuisance covariates—including age, sex, and site—was first established. This configuration was sequentially benchmarked against a full model that integrated the main effect of interest (group). By performing a nested F test to evaluate the incremental reduction in the residual sum of squares between the two models, we derived the empirical statistical significance of the global omnibus group effect.
*(ii) Conducting post hoc pairwise comparisons:* Upon confirming a significant global omnibus group effect, we extracted pairwise linear contrasts directly from the aforementioned full model configuration to pinpoint the specific pairs of groups driving the metabolic divergence. This contrast-based approach permitted all pairwise comparisons to completely share the pooled residual degrees of freedom from the entire sample, thereby suppressing the interference of covariates while more robustly deriving the empirical t-statistics for the adjusted group-specific regression estimate differences.

Furthermore, for patient-specific clinical characteristics such as the disease duration, HOMA2IR, and HOMA2B, independent two-group linear comparisons between biotype 1 and biotype 2 were performed while adjusting for the same confounding factors. All multiple comparisons were corrected using the FDR procedure. Specifically, FDR correction was applied first to the global omnibus group effects across all clinical indicators, and subsequently to the entire pool of post hoc pairwise contrasts and patient-specific two-group comparisons.

### Statistical comparison of behavioral phenotypes across biotypes

To explore whether the neurophysiological biotypes exhibited distinct cognitive impairment patterns and to systematically characterize the cognitive variations across biotypes and HCs, we conducted intergroup comparisons of their cognitive functions at the individual level. Given that comprehensive clinical cognitive metrics were exclusively available within the UKB database, all behavioural intergroup comparison analyses in this section were strictly restricted to the UKB participant subsample. A total of 7 core cognitive indicators tapping into diverse behavioural domains, including trail-making #1, trail-making #2, symbol digit substitution, numeric memory, pairs matching, fluid intelligence, and matrix pattern completion, were integrated. Prior to statistical modelling, the signs of the cognitive scores possessing inverse clinical meanings were mathematically inverted to guarantee a uniform directional interpretation across all the behavioural axes, where higher values universally denoted superior cognitive performance.

We evaluated the behavioural differences across groups using the identical statistical framework established for the clinical and metabolic profiles. Specifically, we applied the aforementioned ANCOVA model and subsequent contrast-based post hoc comparisons, controlling for age, sex, site, and education level as nuisance covariates. The resulting empirical p values were strictly corrected for multiple comparisons by utilizing the FDR procedure.

### Multivariate brain–phenotype association analysis

We employed a PLS correlation analysis to systematically characterize the multivariate coupling patterns between the whole-brain morphometric deviation maps and multidimensional clinical phenotypes within the different neurophysiological biotypes of T2D patients. As a powerful, data-driven multivariate statistical approach, this method decomposes the cross-covariance matrix between two data blocks to identify optimal latent variables that maximize the spatial association between the two datasets. In this study, the PLS correlation framework was deployed across two independent clinical analysis configurations and executed separately within each biotype:

*(i) Brain deviation–clinical metabolic association analysis:* The clinical metabolic data utilized in this study were derived from 9 indicators shared across all three independent datasets: HbA1c, BMI, disease duration, TC, TG, HDL, LDL, SBP, and DBP. In this PLS correlation configuration, the input brain matrix X was the individualized deviation score matrix across 318 cortical regions for patients belonging to a specific biotype, whereas the phenotype matrix Y_1_ contained the participants’ values for the aforementioned 9 clinical metabolic indicators.
*(ii) Brain deviation cognitive association analysis:* In this PLS correlation configuration, the input brain deviation score matrix X was subset to include only the UKB participants within a specific biotype, while the corresponding behavioural matrix Y_2_ integrated 7 core cognitive performance metrics provided by the UKB database. Prior to conducting statistical modelling, the signs of the cognitive scores possessing inverse clinical meanings were mathematically inverted to guarantee a uniform directional interpretation across all the behavioural axes, where higher values universally denoted superior cognitive performance.

The PLS correlation analysis decomposed the relationships between the brain deviations and phenotypes into an orthogonal set of latent variables that maximized the cross-block covariance. These latent variables were linear combinations of the original variables from both datasets, consisting of singular vectors and singular values. Prior to conducting the analysis, all columns in the X and Y matrices were subjected to standard Z score normalization. The cross-covariance matrix between the two data blocks was subsequently computed as R = X^T^Y and subjected to singular value decomposition (SVD):

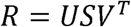

The diagonal elements of the singular value matrix S determined the proportion of cross-block covariance explained by each individual latent variable. The left singular vectors U and right singular vectors V represent the feature weights for the brain regions and phenotypic indicators, respectively, reflecting their contributions to the latent variables. The i-th latent variable comprises the i-th left singular vector, the i-th right singular vector, and the i-th singular value. The i-th singular value specifies the cross-block covariance captured between X and Y by the corresponding i-th latent variable. By projecting the original standardized data matrices onto their respective feature weight vectors, brain scores and phenotype scores were derived for each participant.

To evaluate the statistical significance of each latent variable, we conducted a nonparametric permutation test by randomly shuffling the subject rows 10,000 times. For each individual permutation trial, the cross-covariance matrix was recalculated and subjected to SVD. Consequently, we constructed empirical null distributions for both the singular values and the variance explained of the latent variables. The empirical statistical significance of each latent variable was estimated by benchmarking the observed empirical values against their respective permutation null distributions. Furthermore, to pinpoint specific cortical regions or phenotypic variables that stably contributed to the significant latent variables, a bootstrap resampling procedure with 1,000 iterations was executed with replacement at the subject level to estimate the robustness of the weight coefficients. To overcome the mathematically inherent sign indeterminacy of singular value decomposition, a sign alignment protocol was enforced for the weight vectors across all bootstrap trials relative to the reference weights. The bootstrap ratio (BSR) for each individual feature was quantified by dividing its original weight by its bootstrapped standard error. Features with |BSR| > 2 were identified as core, stably contributing elements that possessed a significant and stable contribution to the observed multivariate coupling patterns.

### Imaging–transcriptomics spatial association analysis

To elucidate the potential molecular mechanisms underlying the individualized brain morphometric abnormality patterns observed within different T2D biotypes, we conducted a multivariate PLS regression analysis to spatially couple the macroscale cortical deviation maps with the microscale postmortem gene expression profiles derived from the AHBA ^35^. The raw microarray expression data from the 6 postmortem human donors provided by the AHBA were normalized and preprocessed using the official abagen toolbox (https://github.com/rmarkello/abagen) ^69^. Given that tissue blocks covered the right hemisphere in only two of the six donor brains, our imaging–transcriptomics mapping was strictly restricted to the left hemisphere to safeguard the continuity and reliability of the spatial transcriptomic data. This process ultimately yielded a spatial gene expression profile across 156 cortical regions encompassing 8,470 genes ^30^.

In the PLS regression modelling framework, the gene expression matrix served as the predictor variable, whereas the biotype-specific average cortical deviation map served as the response variable. The algorithm decomposed the spatial relationships between the brain deviations and the transcriptome into a set of orthogonal latent components. We quantified the proportion of variance explained in the cortical deviation pattern by each latent component and further assessed the spatial alignment between the derived gene scores and the empirical anatomical deviation map using the Pearson correlation coefficient. The spin test ^42^ (10,000 iterations) was subsequently used to test the statistical significance of each latent component. By benchmarking the observed empirical variance explained and the spatial correlation coefficients against their respective 10,000 spin null distributions, we derived a p value for each latent component. For the core latent components demonstrating significant spatial coupling, a region-level bootstrapping protocol with 5,000 iterations was implemented to evaluate the reliability of the individual gene contributions to the multivariate association pattern. To overcome the mathematically inherent sign indeterminacy of singular value decomposition, a sign alignment scheme was enforced for the weight vectors across all bootstrap trials relative to the reference weights. The final stability of the contribution of each gene was quantified using the bootstrap ratio (z score), which was calculated by dividing its original weight by its bootstrap standard error estimated from the 5,000 resampling iterations, representing the ultimate reliability of the biological contribution of that gene.

To further explore the molecular underpinnings and biological pathways associated with the distinct patterns of brain deviations, we generated genome-wide rank-ordered gene lists in both descending and ascending orders, utilizing the bootstrap z scores of the genes as the ranking metric. Subsequently, GSEA ^36^ was performed across the KEGG pathways and the three core domains of GO utilizing the clusterProfiler package ^70^. Statistical significance was rigorously defined as FDR-corrected *p*-value < 0.05.

### Cell-type enrichment analysis

To elucidate whether the genes associated with brain deviation patterns exhibited preferential expression within specific cell types, we incorporated seven canonical cell classes that were previously defined by Seidlitz et al.^44^ via a meta-analysis of five independent single-cell sequencing studies. These cell types included microglia, endothelial cells, oligodendrocyte precursor cells, oligodendrocytes, astrocytes, excitatory neurons, and inhibitory neurons. We subsequently evaluated the observed median ranks of the marker genes for each cell type within the PLS regression-derived, genome-wide rank-ordered list. To assess the statistical significance of these rankings, a nonparametric permutation testing framework (10,000 iterations) was implemented to generate empirical null distributions and calculate empirical *p* value (*p*_perm_) for each cell type. Crucially, all *p*_perm_ values were adjusted for multiple comparisons using the FDR method, with an FDR-corrected *p* value < 0.05 indicating significant cell type-specific enrichment.

### Sensitivity analyses

To validate the robustness and generalizability of the identified T2D neurophysiological biotypes, we conducted four distinct sensitivity analyses. In each analysis, upon re-executing the biotyping pipeline under specific modified conditions, we evaluated the consistency of the biotypes by assessing either the spatial correlations between the newly derived and original group-level deviation patterns or the individual-level concordance among the biotype assignments. The specific analysis details are as follows:

*(i) Leave-one-site-out analysis*: To mitigate the potential biases driven by specific imaging scanners, one of the six independent imaging sites was iteratively excluded, and k-means clustering was reapplied to the remaining samples.
*(ii) Cross-cohort generalizability*: To assess the stability of the biotype structures across different populations, the full sample was explicitly partitioned into two independent cohorts (the Chinese cohort and the UKB cohort), and the clustering algorithm was independently re-executed within each cohort.
*(iii) Robustness to site effect estimation strategies*: To verify that the biotyping results were robust to the strategy used for estimating site effects, we estimated an alternative deviation matrix by reconstructing the normative models (Fig. S6a). Specifically, the HCs within each site across the three datasets were stratified by sex and sorted in ascending order of age. A deterministic split-half strategy was then implemented by assigning the odd-indexed rows to the training set (HCs_train) and the even-indexed rows to the testing set (HCs_test). This stratified alternating sampling approach guaranteed nearly identical age and sex distributions between the two subsets within each site. The pooled HCs_train across all sites was merged with the reference lifespan normative cohort ^30^ to reconstruct and retrain the GAMLSS models, thereby absorbing site-specific variance. The independent testing set, comprising the pooled HCs_test set and all patients with T2D, was then projected onto the retrained curves to estimate individual quantile scores. These quantiles were subsequently transformed into standard Gaussian-distributed Z scores using quantile-randomized residuals based on the fitted Johnson’s Su distribution. The detailed procedures for model training and deviation score calculation are described in our previous study ^30^. Finally, the identical clustering strategy was applied to this alternative deviation matrix.
*(iv) Alternative cortical parcellation:* To ensure that the identified biotypes were not driven by the specific spatial resolution or boundary definitions of the chosen brain atlas, we replicated the entire analytical framework using an alternative parcellation scheme. Specifically, the regional morphological features were re-extracted based on the DK-219 atlas. The individualized deviation scores were re-estimated for these 219 regions, and the identical clustering procedure was applied to the newly generated deviation matrix.

## Data availability

The original raw data are not publicly available due to data sharing agreements and privacy regulations. The UK Biobank data are available to researchers upon approved application via https://www.ukbiobank.ac.uk/enable-your-research/register. Data used in this study are available in the UK Biobank under application number 415382. The Allen Human Brain Atlas is available at https://human.brain-map.org/static/download. The canonical cell-type data can be downloaded from https://github.com/jms290/PolySyn_MSNs/tree/master/Data.

## Code availability

The codes for this study are available on GitHub (https://github.com/wangxyue/T2DM_Neurophysiological_Biotypes). The software packages used in this manuscript include MRIQC v0.15.0 (https://github.com/nipreps/mriqc), QuNex v0.93.2 (https://gitlab.qunex.yale.edu/), HCP pipeline v4.4.0-rc-MOD-e7a6af9 (https://github.com/Washington-University/HCPpipelines/releases), FreeSurfer v6.0.0 (https://surfer.nmr.mgh.harvard.edu/), FSL v6.0.5 (https://fsl.fmrib.ox.ac.uk/fsl/fslwiki/), Connectome Workbench v1.5.0 (https://www.humanconnectome.org/ software/connectome-workbench/), MSM v3.0 (https://github.com/ ecr05/MSM_HOCR/), Python v3.13 (https://www.python.org), MIND (https://github.com/isebenius/MIND), R v4.5.3 (https://www.rproject.org), GAMLSS package v5.4-3 (https://www.gamlss.com/), NbClust package v3.0.1 (https://www.rdocumentation.org/packages/NbClust/versions/3.0.1/topics/NbClust), neuromaps toolbox v0.0.5 (https://github.com/netneurolab/neuromaps), MATLAB R2019b (https://www.mathworks.com/products/matlab.html), abagen toolbox v0.1.3 (https://github.com/rmarkello/abagen), clusterProfiler v4.18.4 (https://bioconductor.org/packages/release/bioc/html/clusterProfiler.html), BrainNet Viewer toolbox v20170403 (https://www.nitrc.org/projects/bnv), ggplot2 package v4.0.3 (https://ggplot2.tidyverse.org/). The code for calculating deviation scores is available at https://github.com/brainchart/Lifespan.

## Acknowledgements

This study was supported by the National Natural Science Foundation of China (82021004 and 82327807 to Y.H.; 824B2051 to X.Y.L.; T24B2012 to L.L.S.; 82360343 to L.P.Z.; 82330058 and T2341014 to S.J.Q.), and the Beijing Natural Science Foundation (JQ23033 to M.R.X.). This study used the UK Biobank Resource under application number 415382. We want to thank all the participants and researchers from the UK Biobank, as well as the GZUCM and GPH cohorts. We also extend our gratitude to Weiwei Wang, Minhui Zhu, Cheng Zhu, and Maolin Li for their valuable assistance in the manual visual inspection and quality control of the structural MRI data.

## Author contributions

X.Y.W., L.P.Z., S.J.Q., and Y.H. conceptualized the study. Y.H. supervised the project. X.Y.W., Z.Z.S., J.C.C., J.T., S.J.Q., and L.P.Z. collected the data for this study. X.Y.W., L.L.S. and X.Y.L. performed data preprocessing. X.Y.L. provided the established normative model as well as the cortical morphometric networks for the validation subsets, and contributed to the methodological implementation. X.Y.W. conducted the formal data analysis, visualization, and validation. Z.Z.S., J.C.C. and M.R.X provided analytical suggestions. X.Y.W., X.Y.L., M.R.X., and Y.H. interpreted the data and refined the analysis. X.Y.W. and Y.H. wrote the original manuscript draft. X.Y.W., X.Y.L., Y.H., J.C.C., Z.Z.S., L.P.Z., and S.J.Q. reviewed and edited the manuscript. All authors reviewed the final manuscript.

